# Tracing 3D apoplasmic complexity in vascular tissue of live conifer needles by X-ray computed tomography

**DOI:** 10.1101/2022.01.22.477321

**Authors:** Chen Gao, Sean J. V. Marker, Carsten Gundlach, Henning F. Poulsen, Tomas Bohr, Alexander Schulz

## Abstract

Architecture and conducting area of vascular elements along conifer needles are fundamentally different from broad leaves. We hypothesised that the needles’ unique transfusion tissue offers different mechanisms for water allocation and used multimodal imaging to dissect the critical water exchange interfaces in this xerophytic leaf type. Our study examined intact conifer needles with X-ray computed tomography (μXCT) and water-soluble tracers, allowing to render the functional 3D structure of the water-filled apoplast and the complementary symplasmic domain inside the bundle sheath. Segmentation of these data, together with fluorescence and electron microscopy of axial phloem and xylem elements along the needle, enabled quantification of the dimensions of the conducting tissue complex. The transfusion tracheid system between the endodermis-type bundle sheath and the axial venation formed a sponge-like apoplast domain. Transfusion parenchyma cell chains bridged this domain not directly but as tortuous symplasmic pathways between bundle sheath and axial phloem, which is nearly exclusively accessible at flanks. The transfusion tissue extends the plasma membrane surface for phloem loading and provides a large volume space. We discuss that this unique tissue plays an important role in the subtle interplay between water uptake/storage and sugar transport that has evolved to cope with desiccation stress.

## Introduction

Conifer needles contain a unique tissue in which the axial vascular elements of xylem and phloem are imbedded. This *transfusion* tissue fills a considerable part of the stele and consists of dead *transfusion tracheids* and living *transfusion parenchyma* cells arranged in a complex structure bounded by the bundle sheath. It carries out the complicated task of transporting water – for transpiration and photosynthesis – outward from the xylem, and sugars – produced by photosynthesis in the mesophyll – inward to the phloem. The details of this two-way traffic are not known, in particular since gymnosperms have an endodermis-like bundle sheath (for simplicity “endodermis” from now on) with a Casparian strip (Canny, 1993; Liesche *et al*., 2011). Water transport can take place only transcellularly (Barberon & Geldner, 2014), i.e., via the lumen of the endodermis cells, where it meets sugars on the way to the phloem.

Interestingly, a tissue like the transfusion tissue is not found in angiosperms, not even in grasses, which also have a linear venation structure. There, the vascular bundles are more numerous and there are typically only few cells between the vascular element and the bundle sheath. Moreover, angiosperms leaves are characterised by having different vein classes, even those with needle-like leaves such as rosemary and dill, indicating a division of labour between leaf import–export and nutrient loading–unloading, respectively. It is obvious that the basic morphology and venation pattern in leaves have large influence on the transport efficiency of the vascular system and their ecological adaptations (McAdam *et al*., 2017; Carvalho *et al*., 2018; Baird *et al*., 2021). It is obvious that the unique needle vasculature reflects an ecophyisological demand.

Plant anatomists were fascinated by the special architecture of conifer needles since the late 19^th^ century and have tried to understand the underlying physiology (Strasburger, 1891; Münch, 1930; Huber, 1947; Heimerdinger, 1951). While the xylem hydraulics of pine needles can be predicted in a single-veined leaf model (Zwieniecki *et al*., 2006), reaching understanding of the opposite flow of assimilates turned out to be more challenging in long needles. The challenge was that the tip region of long needles might not contribute to the sugar export if this export is driven by the generally accepted mechanism, the osmotically generated pressure flow (Münch, 1930). When mathematically approached using established models (Jensen *et al*., 2016), it was shown in (Rademaker *et al*., 2017) that for a constant sugar concentration profile in the phloem, there would be a stagnant flow near the tip of needles with a length beyond 50-60 mm, i.e. no sugar export at all from the tip region. While most conifer species have needles within this range, some species, like *P. pinaster*, have needles which are much longer. Universality of Münch transport in the phloem was otherwise predicted to scale well with 70 angiosperm and gymnosperm tree species (Jensen *et al*., 2012) and transport of phloem tracers was seen in needles of up to 450 mm in length (Ronellenfitsch *et al*., 2015; Han *et al*., 2019). A recent paper suggests that long needles avoid a stagnant flow in the tip by structuring the sieve elements of the phloem in a special way. It has been observed that sieve elements come in groups starting from the tip with each new group appearing at the flank of the phloem (Liesche *et al*., 2021). Interestingly, it seems to be only at the flanks the phloem can be loaded with sugar as fibre cells appear to block other possible entry points in many conifers. Combining these observations into a model, hypothesizing that a group of sieve elements only can be loaded with sugar when it is at the flank and that different groups are separated from each other by Strasburger cell rays, it was shown that long needles in this way avoid having a stagnant flow near the tip (Liesche *et al*., 2021).

Needles are textbook examples for xerophytic adaptations minimizing evaporation by a small surface-to-volume ratio and sunken stomata, where high humidity in the stomatal crypt reduces loss of water vapour from the intercellular air spaces of the mesophyll. As the example of sunken stomata shows, special structures indicate functional adaptations and divergence from the biological norm. The models predicting xylem and phloem transport in needles did not take the unique transfusion tissue into account, and thus fail to describe their potential role in vascular transport (Zwieniecki *et al*., 2006; Charra-Vaskou & Mayr, 2011; Liesche *et al*., 2021). Moreover, there is also a lack of investigations into the possible link between the two transport processes in conifer needles as otherwise was done for angiosperm leaves (Barbour & Farquhar, 2004; Buckley *et al*., 2015; Sakurai & Miklavcic, 2021). However, one pioneering study used fluorescent and radioactive tracers to characterise the physiology of the transfusion tissue in pine needles (Canny, 1993). He demonstrated with pulse-chase experiments that xylem mobile dyes and ^14^C-labelled aspartate move from the axial xylem into the transfusion tracheids and with time appears also in the transfusion parenchyma. A role in nutrient exchange was confirmed by micro-particle induced X-ray emission maps showing enrichment of zinc in the transfusion parenchyma (Pongrac *et al*., 2019). Another physiological aspect is that transfusion tracheids of *Taxus baccata* seemed to contribute to desiccation tolerance by reversible deformation (Zhang *et al*., 2014). Finally, the transfusion tissue was discussed to be involved in the symplasmic mode of phloem loading in gymnosperms (Schulz, 2015), suggested from photoactivation experiments of plants with different phloem loading strategies (Liesche & Schulz, 2012).

Since the transfusion tissue separates the axial phloem and xylem transport from the photosynthetic tissue, we hypothesise that it contributes to the allocation of water to the changing demands in transpiration, photosynthesis, and phloem loading. For studying the key parameters of its functionality, we combined fluorescence and electron microscopy of the long needles of the Mediterranean *Pinus pinaster* with live imaging by high-resolution micro-X-ray computed tomography (μXCT). Microscopy allowed us to quantify the conductive area of xylem and phloem along the needle. Computed tomography allowed rendering the fascinating and surprisingly complex 3D structure of the transfusion tissue and calculate the membrane surfaces available for water and nutrient exchange. Our results provide further evidence that it is only the flank of the phloem which can be loaded with sugar. We will also discuss the significance of our data for water allocation and assimilate loading and hopefully pave the way for a deeper understanding of the functionality of the stelar tissue in conifer needles.

## Material and Methods

### Plant material

Needles from *Pinus pinaster* Aiton were collected from a 4-year old tree grown in a greenhouse at the University of Copenhagen, Denmark with16 h: 8 h, day: night at 18°C: 8°C, day: night in winter, and 25°C: 15°C, day: night in summer.

### Water uptake

A small branch of *P. pinaster* was cut from the tree and immersed in water. Needles were excised from the branch with a razor blade under water. Ten needles each were collected and immediately transferred into 2 mL Eppendorf tubes filled with 1.5 ml water and the open end covered with parafilm to prevent evaporation. The tubes were separated into two groups, one group was under light, the other group was under dark. The weight of each tube (including water and needles) was measured every 2 hours using an electronic balance.

### Sample preparation for microscopy

Needles of *P. pinaster* were collected, the outer wall layers of the epidermis gently removed with sandpaper and cut into 10 mm-segments at the positions shown in the top of Fig. S2. The segments were immediately immersed into Karnovsky’s fixative (4% [w/v] paraformaldehyde and 5% [w/v] glutaraldehyde in 0.1M sodium cacodylate buffer, PH 7.4), degassed under vacuum for 1 hour and further fixed for 18 hours at 4°C. After washing in 0.1M sodium cacodylate buffer, the samples postfixed 1% [w/v] osmium in the same buffer for 4 hours at room temperature on a rotator. Further steps, including washing, dehydration in acetone and embedding in Spurr’s resin (Merck Life Science A/S, Søborg, Denmark) followed the standard procedure (Hunziker & Schulz, 2019). To avoid preparation artefacts, 3-mm ends of each segment were discarded. The samples were polymerized in moulds at 70°C for 72 hours and trimmed for electron and fluorescence microscopy.

Ultrathin sections of 70 nm thickness were cut with a diamond knife and ultramicrotome (EM UC7; Leica Microsystems, Wetzlar, Germany). The sections were transferred to film-coated single-slot grids (FCF2010-CU, Electron Microscopy Sciences; Hatfield, PA 19440, USA) and post-contrasted with uranyl-less solution for 3 minutes and lead citrate solution for 3 minutes with thorough washing after each step. For transmission electron microscopy (TEM) a Philips CM 100 was used at 80 kV acceleration voltage. Diameters of sieve elements and xylem tracheids were measured using ImageJ (https://imagej.net/software/fiji/).

Semithin sections of 2 μm thickness were also cut with ultramicrotome and diamond knife. The sections were transferred onto drops of distilled water placed on a microscope slide, dried on a heating plate and stained with 3% (w/v) Coriphosphine O (TCI Europe, Zwijndrecht, Belgium CAS 5409-37-0) for 5 minutes. Images were taken with a wide-field fluorescence microscopy (Nikon Eclipse 80i, Amsterdam, Netherlands) at 450-490 nm excitation and 520 nm long-pass emission filters.

### X-ray μCT

Small branches of *P. pinaster* were cut from the tree on the day before imaging. To avoid xylem embolism, the branch was immersed in water and the needles were excised under water at the base with a razor blade. Needles were immediately transferred to 2 ml Eppendorf tubes with Milli-Q water or contrasting agent solutions where they stayed up to imaging. Imaging was done with an Xradia 410 Versa X-ray microscope (Zeiss, Oberkochen, Germany capable of micro computed tomography (μXCT) and 40-60 kV and 4 or 10x zoom optics. A chamber was designed to stably hold needles during rotation while immersed in the Eppendorf tube (Fig. S1). Reconstructions were performed using the software of the instrument. The results were z-stacks of needle cross sections with the thickness of the effective pixel resolution, chosen between 2.2 and 3.5 μm. The stacks consisted of 900 to 990 slices and were saved in Zeiss’ proprietary txm-format.

As contrast agents, potassium iodide or iohexol (TCI Europe, Zwijndrecht, Belgium; CAS 66108-95-0) (Pratt & Jacobsen, 2018)) were supplied. Potassium iodide and iohexol, dissolved in Milli-Q water to 2.4 mM and 150 mM, respectively, were filtered through 200 nm membrane filters. Incubation started 12 h before imaging. Where necessary for direct comparison, we used custom-made two-chambered tubes for control and tracer, respectively.

### Imaging, segmentation and quantification of μXCT data

For imaging and segmentations, image stacks were imported into ImageJ, saved as tagged image format stacks (.tif), and cropped to contain the entire cross section of the needle or the stele, respectively. Whenever the number of sections in a stack was above the amount the computer workstation could handle, the stack was divided into smaller stacks, also using ImageJ’s “Stacks—Crop (3D)” plugin. To study specific planes inside the stele, we used ImageJ’s “Dynamic Reslice” command after defining the respective regions on a cross section with the line tool (straight or free hand). For segmentation, the stack was subjected to the ImageJ plugin “Trainable Weka segmentation 3D” or to the open source software “Ilastik” (https://www.ilastik.org/documentation/). Three classes of x-ray contrast values were used for training: iohexol-positive tissue, iohexol-negative tissue, and air within and around the sample. The respective areas were chosen with the freehand selection tool on different slices from a stack. The plugin uses these selections to train parameters that fulfil the mean intensity and variance values in the entire stack with these settings: minimum and maximum sigma 1.0 and 8.0, respectively, FastRandomForest, Balance classes, and 33% result overlay opacity of 33 %. Output files were probability maps revealing the three classes in three different channels that cut be visualised in ImageJ’s 3D render plugins or directly imported into the open-source software project FluoRender (https://www.sci.utah.edu/software/fluorender.html). This software enabled us to select aspects of the sample in any orientation and of any slice thickness and make 3D animation videos. Individual channels were visualised in grey values or combined in appropriate colour combinations to show maximum projections or adjust transparency through the selected slice to get an impression of depth (for examples see Fig. 4 and supplemental videos).

Quantification of the surfaces and volumes of the different needle tissues was based on the segmented data at five equidistant z-positions along the scanned middle segment (cf. Fig.1c), which had an axial length of 3.3 mm and 2.1 mm at 3.4 μm and 2.2 μm resolution, respectively. The field of view of the imaging system did only cover part of the needle cross section at 2.2μm resolution. Quantification was done on transverse slices by outlining the respective tissue borders (epidermis+hypodermal fibres, mesophyll, outside endodermis, inside endodermis) and using the ImageJ tool “Analyze—3D-Objects Counter”. This resulted in surface and volume values for whole needle, mesophyll+endodermis+stele, endodermis+stele, and stele, respectively, from which the respective values were derived by subtraction (see Table 1).

**Fig. 1.**
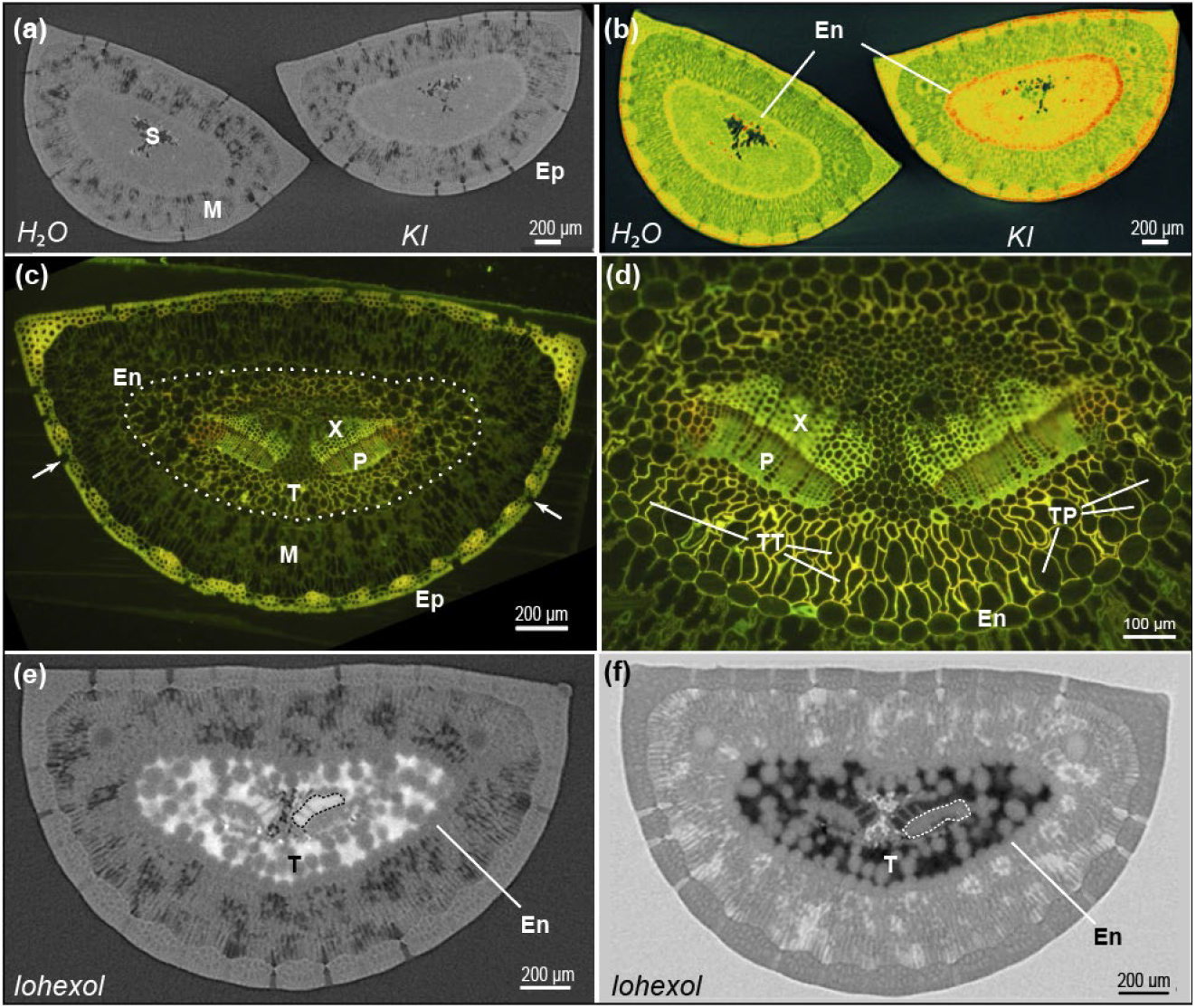
*P. pinaster* needle cross sections from μXCT and fluorescence microscopy. (a) and (b) Sections reconstructed from 3200 μXCT projections with 3.4μm voxel resolution. (a) Single slice of a needle pair, supplied with water and potassium iodide, respectively. Difference in X-ray contrast reveals epidermis (Ep), mesophyll (M) and vascular stele (S). (b) Same pair as (a) but rendered as a 50μm slice with a rainbow look-up table to emphasise the KI accumulation in stele and endodermis (En) as well as mesophyll and epidermis. Note: colour-vision impaired readers are referred to Fig. S5. (c) Needle cross sections stained with the fluorescent dye Coriphosphine O at mid position of the needle (see Fig.S1). The stele contains two vascular bundles with xylem (X) and phloem (P). Epidermis (Ep) with sunken stomata (arrows) borders hypodermal fibres and the mesophyll (M). The endodermis-type bundle sheath (En) is marked with a stippled line (d) Enlarged view of the stele. The transfusion tissue inside the endodermis contains transfusion tracheids (TT) and transfusion parenchyma (TP), both abutting the endodermis. (e) Single slice of a needle incubated in the xylem tracer iohexol. The tracer is only seen in the axial xylem (dotted outline) and the transfusion tracheid system inside the endodermis (En). (f) Another slice from the same μXCT stack but now shown with inverted contrast. Iohexol-positive tissue is black, axial phloem (dotted outline) and the transfusion parenchyma inside the endodermis (En) are grey. T, transfusion tissue.

**Table 1.** Membrane areas available for osmosis and volumes of tracer-positive transfusion tracheids (TT), tracer-negative transfusion parenchyma (TP) and air spaces per mm needle, based on TEM sections and μXCT segmentations, respectively.

| Tissues | Membrane surface |  | Total volume |  | Iohexol positive |  | Iohexol negative |  | Air |  |
| --- | --- | --- | --- | --- | --- | --- | --- | --- | --- | --- |
| | $(mm^2 \cdot mm^{-1})$ | | $(mm^3 \cdot mm^{-1})$ | | | | | | | |
| axial xylem elements <sup>1)</sup> | 6.40 | ±1.45 | 0.018 | ±0.007 |  |  |  |  |  |  |
| axial phloem elements <sup>2)</sup> | 4.01 | ±0.54 | 0.005 | ±0.001 |  |  |  |  |  |  |
| Stele <sup>3)</sup> | 21.35 | ±3.67 | 0.413 | ±0.009 | 0.150 | ±0.008 | 0.252 | ±0.009 | 0.011 | ±0.001 |
| Endodermis |  |  | 0.141 | ±0.014 | 0.002 | ±0.001 | 0.139 | ±0.013 | 0.000 | ±0.000 |
| Mesophyll |  |  | 0.987 | ±0.022 | 0.001 | ±0.001 | 0.865 | ±0.034 | 0.121 | ±0.016 |
| Epidermis incl. hypodermal fibres |  |  | 0.390 | ±0.013 | 0.005 | ±0.003 | 0.371 | ±0.014 | 0.013 | ±0.002 |
| entire needle |  |  | 1.932 | ±0.011 | 0.158 | ±0.009 | 1.628 | ±0.017 | 0.146 | ±0.016 |
| fold increase <sup>4)</sup> | 5.3 |  |  |  | 8.5 |  | 54.5 |  |  |  |
| | | | | | ($\%$) | | | | | |
| Stele |  |  | 21.4 |  | 36.29 | ±2.056 | 61.01 | ±1.958 | 2.70 | ±0.13 |
| Endodermis |  |  | 7.3 |  | 1.06 | ±0.300 | 98.75 | ±0.274 | 0.19 | ±0.19 |
| Mesophyll |  |  | 51.1 |  | 0.13 | ±0.096 | 87.57 | ±2.017 | 12.30 | ±2.11 |
| Epidermis incl. hypodermal fibres |  |  | 20.2 |  | 1.39 | ±1.010 | 95.22 | ±1.092 | 3.39 | ±0.63 |
| entire needle |  |  | 100 |  | 8.18 | ±0.456 | 84.27 | ±0.894 | 7.55 | ±0.96 |
| <sup>1)</sup> Calculated from TEM (circular+rectangular outline)/2); related to tracheid lumina (N=167) |  |  |  |  |  |  |  |  |  |  |
| <sup>2)</sup> Calculated from TEM (rectangular outlines); related to sieve element lumina (N=211) |  |  |  |  |  |  |  |  |  |  |
| <sup>3)</sup> surface of iohexol-positive domains in stele (mean and SD of six segmentations) |  |  |  |  |  |  |  |  |  |  |
| <sup>4)</sup> Extension of the axial phloem's membrane surface by TP, and of axial xylem's and phloem's volume by TT and TP, respectively |  |  |  |  |  |  |  |  |  |  |

## Results

To study the internal structure of intact needles we constructed a cylindrical chamber (Fig. S1) to centre the needles for μXCT imaging, where the needle is rotated 360°. Importantly, each needle was excised from the branch with a sharp razor blade under water, and immediately transferred into a tube containing water or a water-soluble tracer, where it stayed during the x-ray scanning. The dimensions of the scanned image cube depended upon the resolution, for 3.4 μm resolution it was c. 3×3×3 mm within the mid needle segment (see Fig. S2). Fig. 1a shows a single, 3.4 μm thick slice of a control needle on the left side and a needle that was incubated in potassium iodide on the right side. The slice, reconstructed from all μXCT projections, allows to discriminate epidermis, the mesophyll with intercellular air spaces, and the stele. Iodide isotopes were used in biochemical uptake experiments and shown to be mobile in the xylem and membrane permeable (Kiferle *et al*., 2013; Humphrey *et al*., 2019). Here we used iodide because its high atomic number increases X-ray contrast. When rendered as maximum projection of a 50 μm thick slice (Fig. 1b), all tissues of the treated needle showed higher contrast compared with the water control, but particularly in the endodermis. This indicates that iodide moved in the xylem along with the transpiration flow, and laterally spread out into the transfusion tissue up to the endodermis. Accumulation in the endodermis suggests a symplasmic step, i.e., crossing of the inner and outer plasma membrane to finally end in the apoplast of cortex and epidermis (Fig. 1b).

For a detailed analysis and quantification of the different tissues along the needle, we fixed and stained *P. pinaster* needle segments for fluorescence and electron microscopy. Fluorescence micrographs of cross sections (Fig. 1c and d; Fig S2) revealed two vascular bundles with xylem and phloem on the adaxial and abaxial side, respectively: The bundles are embedded in transfusion tissue that is bordered by an endodermis-type endodermis with Casparian strip. It consists of dead transfusion tracheids with lignified walls and living transfusion parenchyma cells, both abutting the endodermis and vascular bundles, respectively (Fig. 1d). Outside the endodermis, mesophyll, hypodermal fibres, and the epidermis follow in concentric layers. As in many xerophytes, stomata are sunken below the epidermal surface under stomatal crypts (Fig. 1c arrows). The general structure is maintained along the entire needle length of 180 mm (see Fig. S2).

Since contrast by potassium iodide only improved the general contrast in X-ray tomography but did not allow further insights into the transfusion tissue, we exploited the similarly iodine-based medical contrast agent iohexol. Iohexol is a water-soluble molecule with a molecular mass of 821 g·mol^-1^, not membrane-permeable but able to pass bordered pits between tracheary elements and, thus, access leaves (Pratt & Jacobsen, 2018; Holmlund *et al*., 2019). Fig. 1e reveals that iohexol strongly enhanced the contrast and was contained by the endodermis to the stelar apoplast, comprising axial xylem and the transfusion tissue, while phloem, transfusion parenchyma and endodermis as well as all tissues outside the endodermis showed the same contrast as water-incubated controls (cf. Fig. 1a: left needle). Fig. 1a and 1e follow the tradition of radiologists with high absorption being brighter than air and iohexol accordingly white. Fig. 1f depicts a similar cross section, now with an inverted look-up table, where high absorption is dark and iohexol therefore black, making it easier to recognise the rounded shape of the grey transfusion parenchyma cells (cf. TP in Fig. 1d).

Since presence of symplasmic bridges – needed for assimilate transport between endodermis and phloem – is not evident in single sections, we explored the 3D volume of a μXCT stack with 2.2 μm resolution and placed a virtual plane along a tangential line inside the endodermis (Fig.2a) and unfolded it in such a way that it became the x-axis, while the axial direction became the y-axis (Fig. 2b). Interestingly, the shape of the transfusion parenchyma cells varied strongly depending on their positions on the flattened sides and flanks, respectively. On both the adaxial and abaxial needle side, the cells had an elongated shape in axial direction (Fig. 2b: sectors labelled 1+5 and 3), while they were clearly isotropic on the flanks (Fig. 2b: sectors labelled 2 and 4). At the adaxial and abaxial sides, the cells also appear to be well connected in both the tangential – and horizontal directions, which is not the case at the flanks, where the connectivity seems much lower in these directions. Instead, the cells at the flanks make radial connections as can be seen in Fig. 3, where we look at radial planes including the flanks of the phloem. The isodiametric transfusion parenchyma cells seem to touch each other, but do not form direct radial cell bridges towards the phloem. To our surprise, we found longitudinal cell chains near the flank of each phloem bundle. Perpendicular planes give evidence that these cell chains are forming a tube-like structure rather than a sheet (see also Fig. S3). The location of these cell chains near the flanks of the phloem is very interesting, and it is obvious to speculate that they, together with the Strasburger cells also found in this area, play an important role in phloem loading (Liesche & Schulz, 2012).

**Fig. 2.**
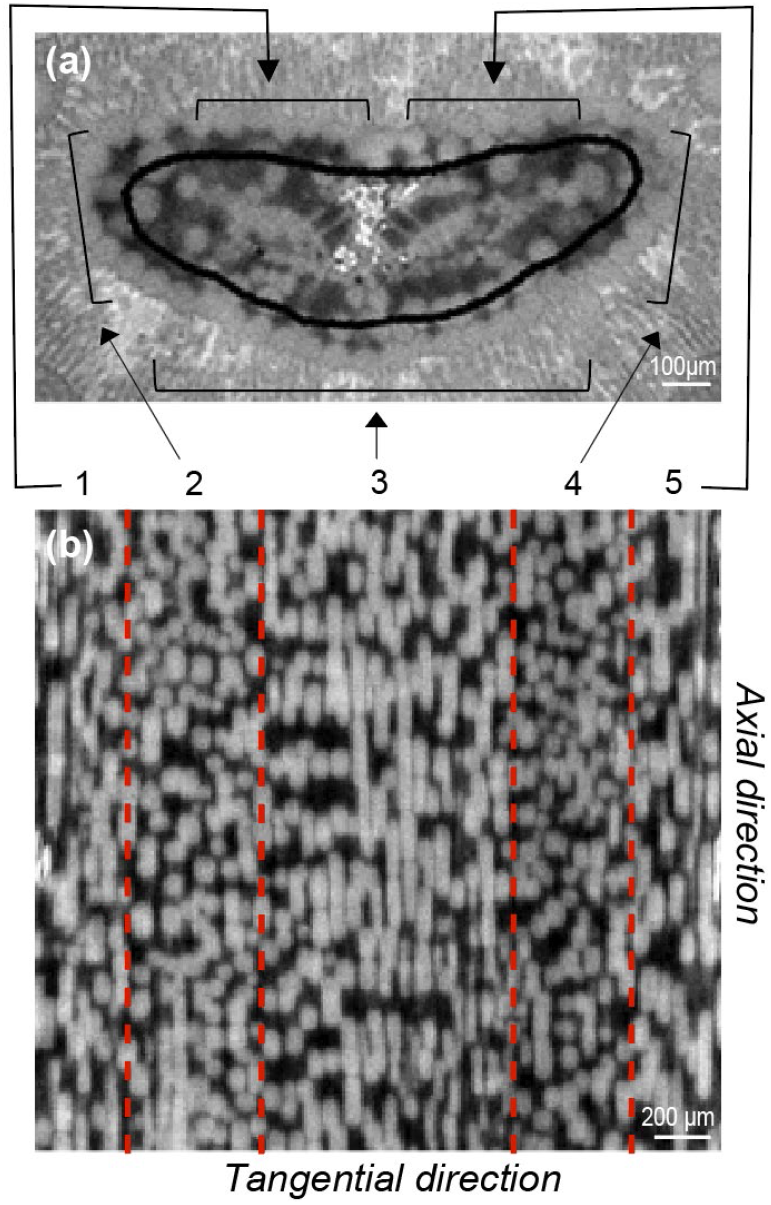
Two aspects of a 3D μXCT stack of a *P. pinaster* needle after iohexol uptake using the same visualisation as Fig. 1f, but with 2.2 μm resolution. (a) Single cross section with kidney-shaped black line indicating the tangential plane just inside the endodermis which is unfolded in (b). (b) Unfolded plane of the entire scanned volume (~2 mm). The shape of the grey transfusion parenchyma cells varies over the unfolded plane as indicated by the dashed red lines. Near the flanks of the bundle sheath, sector 2 and 4, the transfusion parenchyma cells are isotropic and not well connected in either the tangential – or axial directions. By contrast, the transfusion parenchyma cells in sectors 1,3 and 5 are anisotropic, being elongated in the axial direction, and in close contact with each other in axial and tangential direction

**Fig. 3.**
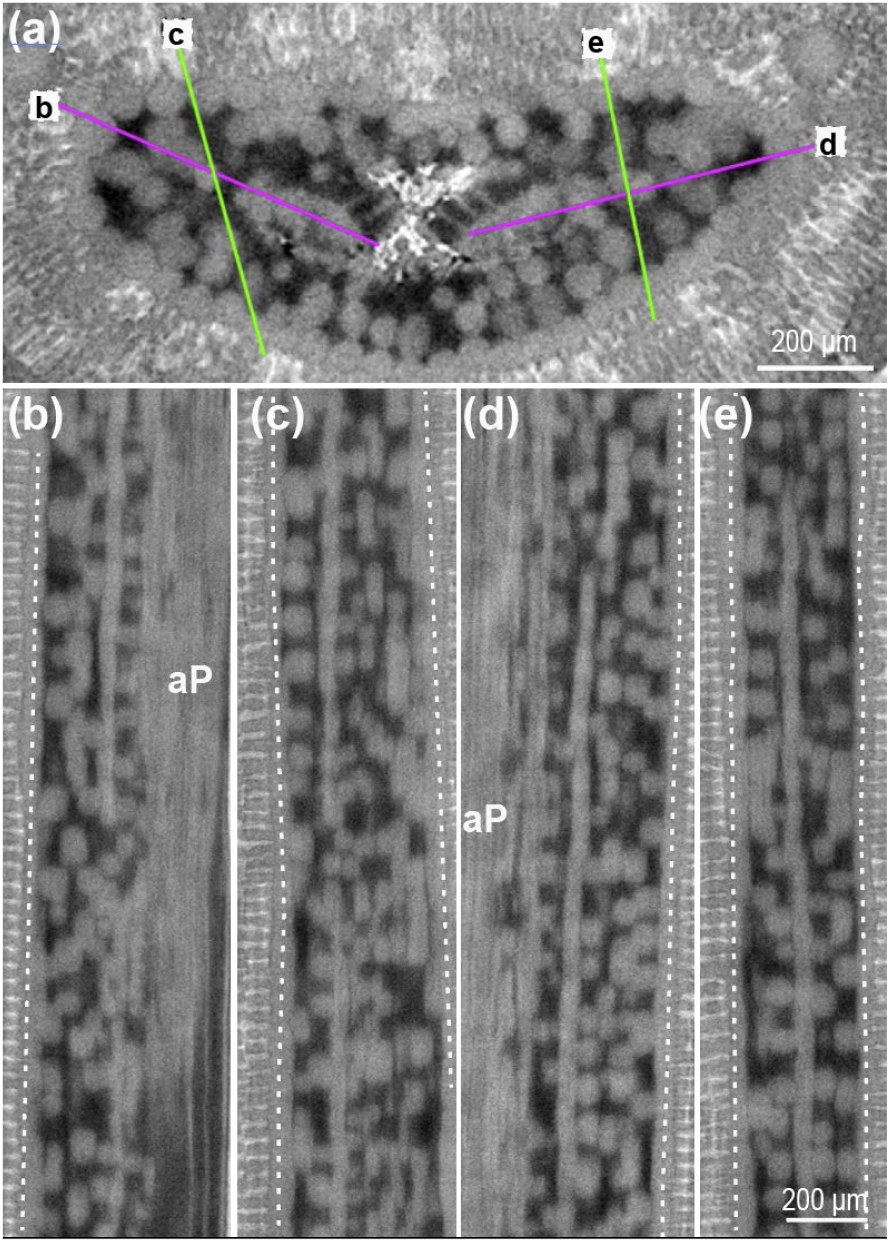
Five 2.2-μm thin slices through the 3D μXCT stack, depicted in Fig. 1f. (a) Orientation of the longitudinal sections shown in the lower panels is indicated, where (b) and (d) include the axial phloem bundles and their outer flanks (magenta), while (c) and (e) run crosswise (green). (b)-(e) While most transfusion parenchyma cells are individual isodiametric cells, there is one cell chain on either flank of the axial phloem (aP) that seems to form a tube-like structure with a diameter of some 40 μm, as shown in view pairs in (b), (c) and (d), (e), respectively. Fig. S3 shows the same tube-like structure as (d) but 3D-rendered after segmentation as a 27 μm slice. Endodermis, stippled lines

**Fig. 4.**
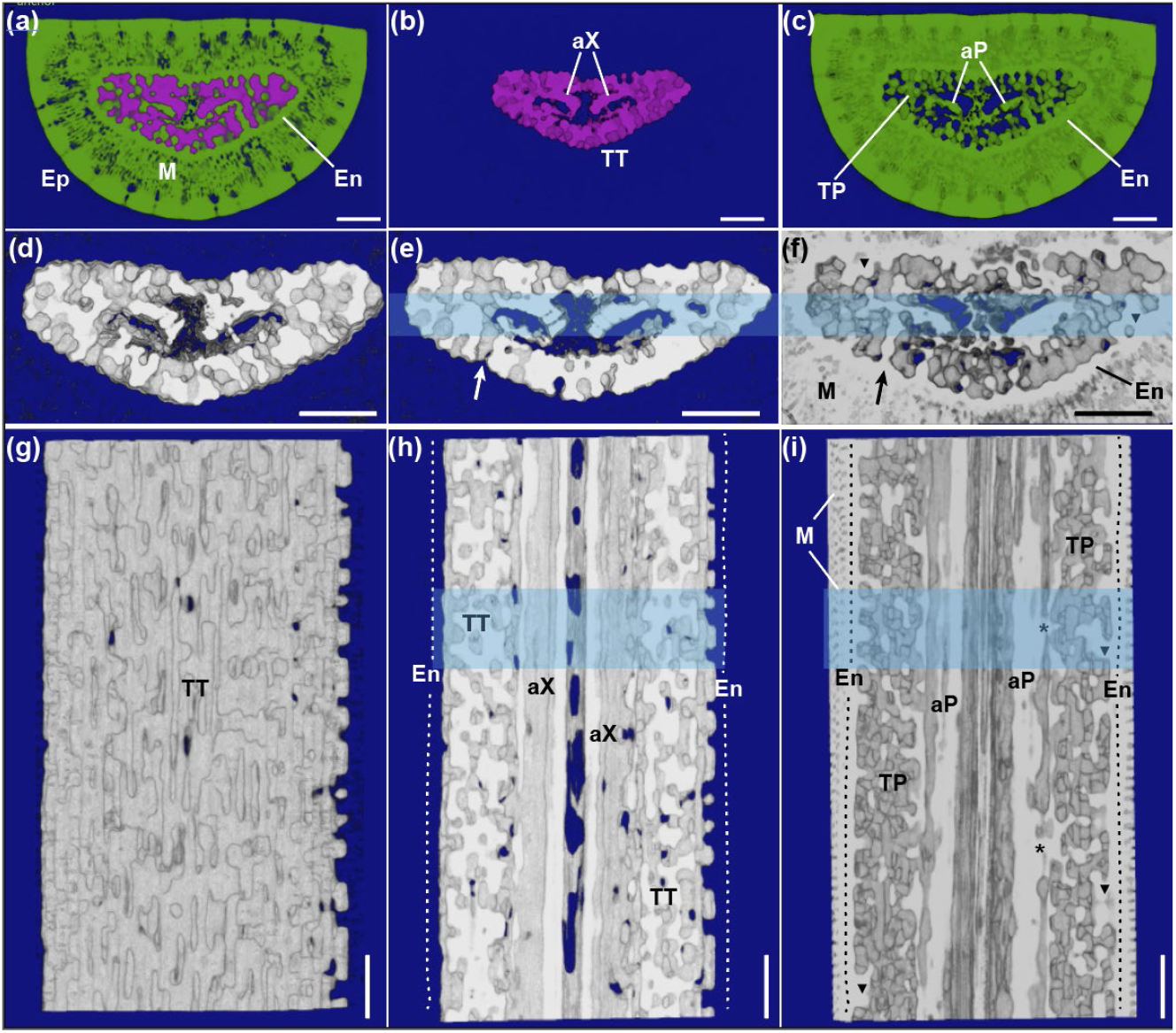
Segmentation and 3D rendered aspects of a needle, imaged by μXCT, in transverse and axial direction. (a)–(c) 3.4μm resolution. (a)-(c) The image stack of a needle incubated in iohexol was rendered as a 50 μm slice and segmented into three classes: iohexol-positive tissue (magenta), iohexol negative tissue (green) and air (blue) showing epidermis (Ep), mesophyll (M) and stele (S). (a) All three classes are seen; (b) only the apoplast inside the stele is depicted with axial xylem (aX) and transfusion tracheids (TT). (c) Only symplasm and cell wall space are shown with axial phloem (aP) and transfusion parenchyma (TP) inside the endodermis (En). (d)–(i) 3D rendered aspects of the stele at 2.2μm resolution. (d) Transfusion tracheid system seen through the entire length (2.1 mm). (h) and (i) 3D rendered 250-μm transverse slice segmented for apoplast (e) and symplasm (f), respectively. Transfusion parenchyma cells bulging out from the endodermis (▾) fill the gaps in the transfusion tracheids system (e.g., at arrow in (e) and (f)). (g) Transfusion tracheids (TT) from a porous, sponge-like system. (k) and (l) 3D rendered 140-μm longitudinal slice segmented for apoplast (h) and symplasm (i), respectively. Axial xylem (aX) and phloem (aP) complement the two bundles and are surrounded by transfusion tracheids and transfusion parenchyma, respectively. Extensions of the phloem (*) flank the axial phloem. There are rarely connections between these and endodermis-associated cells (▾). Location of the rendered longitudinal and transverse slices is light-blue shaded in e, f and h, i, respectively. Scale bars, 250 μm

The clear contrast of iohexol treated needles enabled us to define three x-ray-contrast classes, segment the scanned needle volume and visualise it as 3D rendered structures and animations. Fig. 4 a-c show the same sample as Figs. 2 and 3 as 50 μm thick 3D-rendered slice to visualise the apoplast inside the stele and the entire living tissues of the needle. The iohexol-positive apoplast domain inside the endodermis is shown in magenta, all other tissues in green, and air in blue, respectively. Details of such renderings are shown at higher resolution in Fig. 4d–i. Image processing allowed choosing slices of different thickness in all three dimensions, here visualised as transverse or longitudinal aspects either of the entire scanned volume (Fig. 4d and g) or thinner parts of it (Fig. 4 e, f, h and i). Reconstructions of the iohexol-filled space shows that the apoplast forms a sponge-like porous structure that is well linked with the axial xylem. Comparison of the apoplast structure with the shape and size of transfusion tracheids (see Fig. 1d) indicates that μXCT is not able to resolve single transfusion tracheids, but groups of two to three tracheids. By contrast, chains of transfusion parenchyma cells are clearly resolved extending from the axial phloem and bulging out form the endodermis (Fig. 4f and 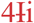; see also supplemental movies of 3D animations).

To quantify the dimensions of axial tracheids and sieve elements along the needle and to assess the total conducting area of xylem and phloem, we counted the number of conducting elements in fluorescence microscopic sections as the ones shown in Fig. 1 and processed cross sections at the same positions for transmission electron microscopy. Examples of the ultrastructure of phloem and xylem are depicted in Fig. 5. Fig. 5a shows the regular arrangement of sieve elements with thick cell walls, a phloem ray consisting of Strasburger cells and only a few phloem parenchyma cells, recognised by the dark-stained tannic acids in their vacuole. Sieve elements were rectangular in their outline and had membranous contents, sieve-element plastids, and mitochondria in an otherwise electron translucent lumen. Wavy walls in the distal sieve elements showed indications of the onset of obliteration. All features were in accordance with the well-known ultrastructure of conifer phloem (Schulz, 1990). Fig. 5b illustrates the xylem consisting of many tracheids with thick, lignified cell walls that were connected by large, conifer-type bordered pits having apertures on either side and a central torus (Schulte & Hacke, 2021). The parenchyma of a xylem ray contained cytoplasm, indicative of being alive when fixed for TEM.

**Fig. 5.**
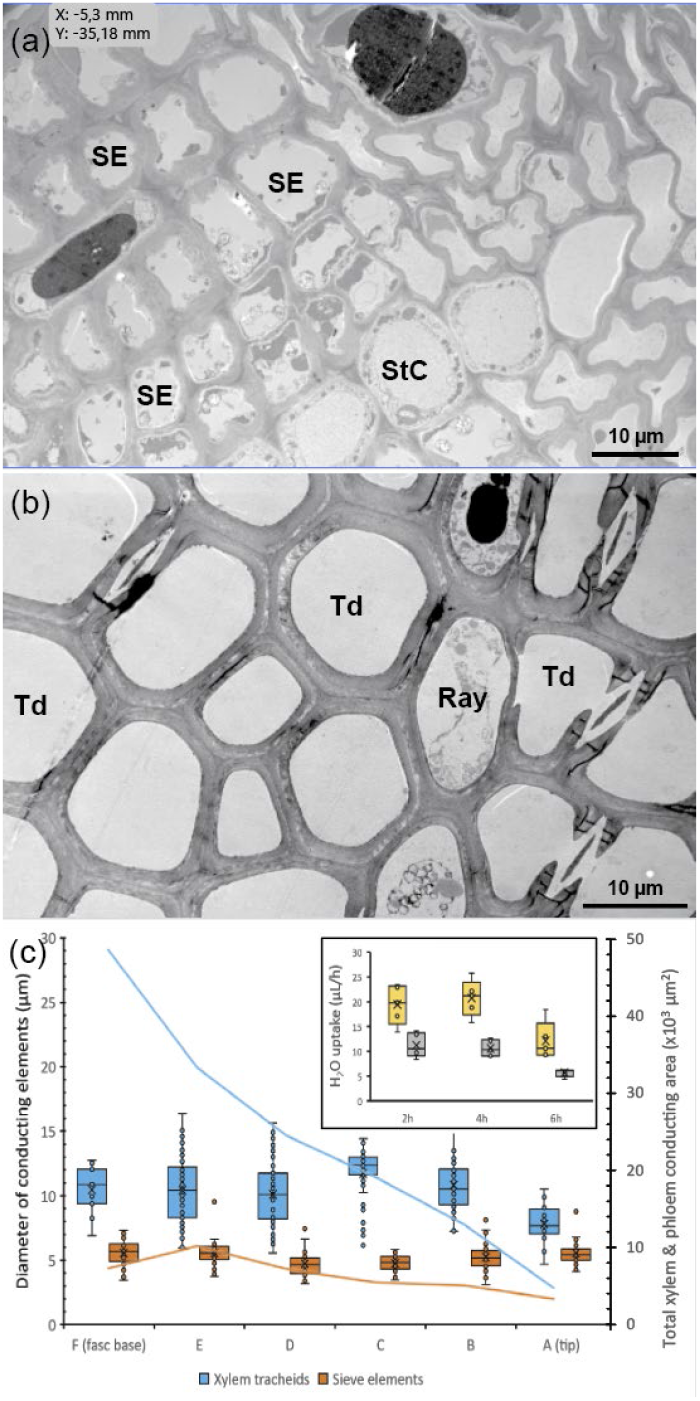
Conducting areas for phloem and xylem transport in needles. (a) and (b) Transmission electron micrographs of the phloem and xylem. (a) Numerous sieve elements (SE) with thick cell walls contain membrane complexes, plastids, and mitochondria and are interspersed by a few parenchyma cells with electron-opaque vacuoles and a phloem ray with four Strasburger cells (StC). (b) Tracheids (Td) of the xylem are characterized by thick, lignified cell walls and an empty lumen. Four bordered pits with apertures on either side and a central torus are depicted. The only cells with cytoplasmic contents are found in the ray. (c) Diameters (box chart) and total conducting areas (lines) of xylem tracheids (blue) and sieve elements (orange) from base to tip starting inside the fascicular sheath (F), see location in Fig. S2. The insert in (c) shows quantification of water uptake per hour after needle detachment in the presence (yellow) and absence of light (gray).

Mature *P. pinaster* needles reach a length of more than 180 mm. We expected that the number of conducting elements and/or the conducting area decrease towards the tip (resp. increase towards the base) reflecting the decreasing transpiratory water loss towards the tip and increasing amount of assimilates produced along the needle. Fig. 5C shows that the diameter of conducting elements was quite constant along the needle except for the tip segment, which contained smaller tracheids than the other segments. By contrast, the conducting area, defined as the number of conducting elements times their cross-sectional area, decreased down to c. 18 % for the axial xylem, and only to 35% for the axial phloem. Interestingly, the conducting area for the xylem dropped strongly from the fascicular base to the next segment, while these segments showed a constant conducting area in the phloem (Fig. 5c).

To assess the possibility that removal of the needle led to a wound response, we quantified the volume of the needle’s functional xylem and tested whether water uptake changed over time. Fig. 5c insert shows that needles over the initial four hours took up some 20 μL and 10 μL per hour in light and dark, respectively. A moderate response to preparation was indicated thereafter. The difference in water uptake in the presence and absence of light indicates that less water is allocated to the mesophyll and transpiration due to the light dependency of stomata opening and/or photosynthesis.

TEM and segmentations allowed to estimate key quantitative parameters of *P. pinaster* needles (Table 1). TEM data were used to assess the conducting volume of axial xylem and phloem, defined as the volume bordered by cell wall and plasma membrane, respectively. Stacks reconstructed from μXCT scan were used to quantify tissue surfaces and volumes and intercellular air spaces. They were estimated to be 21 % stele, 7 % endodermis, 51 % mesophyll and 20 % epidermis with hypodermal fibres. Table 1 indicates that the mesophyll included 12.3 % air. This is certainly an underestimate, since smaller air pockets are not resolved at 3.4 μm resolution. Air in the epidermis is due to the stomatal crypts which are well resolved in the μXCTs. As also seen in Figs. 1 and 4, small air inclusions were found in the centre of the stele of all needles scanned. According to 3D analysis, these are fibre-like axial structures extending throughout the scanned volume (Fig. S4). In addition to these structures, a rupture of the endodermis and air intrusion in the stele was observed in one experiment after long X-ray exposure. Stelar air pockets have been observed before in needle and seedling micrographs using cryo-SEM or μXCT (Roden *et al*., 2009; Bouche *et al*., 2016).

As pointed out above, iohexol can neither pass plasma membranes nor the endodermis because of its molecular size and the Casparian strip between endodermis cells. These properties enabled us to estimate the membrane surface available for osmosis and for other membrane transport processes. Tracer distribution was exactly complementary to the axial phloem and transfusion parenchyma as evident when comparing the image pairs of apoplast and symplasm inside the stele (Fig. 4). Thus, the surface of apoplast domains inside the stele was considered identical to the membrane surface of axial phloem and transfusion parenchyma. Remarkably, the surface area and volumes varied only little at the five positio ns (Table 1). While the membrane surface of all individual sieve elements amounts to 4 mm^2^ per mm needle according to TEM measurements, segmentations of the membrane surface inside the stele – but not resolving the sieve elements – is estimated to c. 21 mm^2^. Accordingly, the phloem-element membrane surface is extended more than fivefold by the transfusion parenchyma (Table 1). Applying the same calculation to the conducting volumes of axial xylem (18 nL) and transfusion tracheid system (150 nL), the extension is more than eightfold, and to that of the axial phloem and the transfusion parenchyma even more than 50-fold (Table 1).

## Discussion

We have studied the transfusion tissue by X-ray tomography of intact needles and supplied them at their base with a water-dissolved medical contrast agent, iohexol, that was taken up and transported into the transfusion tissue. Iohexol did not cross the plasma membranes of the transfusion parenchyma. Thus, we directly obtained maps of transfusion tracheids in high contrast, where water is believed to be transported “out” under negative pressure driven by evaporation, and transfusion parenchyma in low contrast, where the sugars are believed to be transported “in” by diffusion and/or bulk flow driven by osmosis (Schulz, 2015). Transfusion tracheids seem to form a continuum surrounding the transfusion parenchyma. The full complexity of the transfusion tissue became first evident after image analysis and 3D rendering (Fig. 4). It has the appearance of a sponge, where the water moves out through the continuum, and the sugar moves in through the holes, i.e., the transfusion parenchyma, cells which are coupled by plasmodesmata. The interaction between the two cell types takes place via the aquaporins allowing water, but not sugar, to pass (Laur & Hacke, 2014). Independent of their individual shapes, the bulging outline of transfusion parenchyma cells indicates that they are turgescent and thus that the needles are intact and well-watered needles during μXCT.

We did not find direct radial connections between endodermis and phloem that would provide short paths for sugar transport. Instead, in transverse needle sections they appeared as isolated structures consisting of one to a few cells (Fig. 1), even though symplasmic continuity between endodermis and phloem was shown by photoactivation experiments (Liesche & Schulz, 2012). Axial aspects of the needle confirmed Huber’s observation that transfusion parenchyma cells are elongated at the adaxial and abaxial needle sides, but not at the flanks, where the cells appear isotropic (Fig. 2) (Huber, 1947; Heimerdinger, 1951). However, Huber’s idea of a well-ordered structure with one transfusion parenchyma cell pr. endodermis cell was not seen. Instead, the transfusion parenchyma cells connecting to the endodermis appear randomly distributed. It was also seen that transfusion parenchyma cells at the adaxial and abaxial needle sides were elongate and abutted each other in the tangential and axial directions (Fig. 2), very much in contrast to the cells at the flanks, which were isodiametric (Fig. 3). Accordingly, the pre-phloem pathway of photosynthates seems complicated by the fact that it cannot simply move radially inward at the flat needle sides but must move sideways out toward the flanks. The high connectivity in the axial direction might help to avoid a possible bottleneck in the sideways transport and to ensure a steadier flow towards the phloem while still allowing waterflow out through the transfusion tracheids. At the flanks, only around 50 μm outside the phloem, a tube-like cell chain is found at either side of the needle (Fig. 3, Fig. S3), which many of the symplasmic bridges between the endodermis and the phloem seem to go through. This side of the axial phloem is bordered by a 3-cell array of Strasburger cells as already described by Carde (Carde, 1973). These observations provide further evidence that phloem loading only takes place at the flanks, and they are well in line with the model proposed by Liesche et al. (Liesche *et al*., 2021) describing how long conifer needles avoid stagnation in the sugar transport.

In the well-watered situation, the transfusion tissue is obviously involved in both the post-xylem nutrient and the pre-phloem photosynthate transport, as it separates sources and sinks of these processes. Water-dissolved nutrients move in the transfusion tracheid system to the endodermis, cross this layer and are utilised in the mesophyll while the water evaporates from the apoplast space through stomata. This transport takes a combination of three pathways: It can move in the apoplast through the lumen of the transfusion tracheids and permeable cell walls, it can cross the cell membranes through channels and transporters, and it can go cell-to-cell through plasmodesmata (Canny, 1993). A similar combination of pathways is the well-known - but inverted - nutrient uptake pathways in roots, where modelling showed that the radial water conductivity mostly depends on the membrane conductivity of the endodermis (Couvreur *et al*., 2018). In the root as well as in in the needle, water and dissolved nutrients have to enter the symplasm of the endodermis due to the Casparian strip (Mayr *et al*., 2014; Hacke *et al*., 2015). Opposite to the water flow, photoassimilates move symplasmically from the mesophyll through endodermis, transfusion parenchyma and Strasburger cells to the sieve elements of the axial phloem. Accordingly, the opposite flows meet in the cytosol of endodermis cells. Exactly how the endodermis handles this bi-directional transport is not known.

An obvious question is why conifer needles develop a complex transfusion tissue that increases length and complexity of the radial transport pathways considerably. When comparing leaf venations across the plant kingdom, this distance influences the general water hydraulic conductance and, at the same time, photosynthetic efficiency (Brodribb *et al*., 2005; Brodribb *et al*., 2007; McAdam *et al*., 2017). However, the disadvantage arising from a long distance might be less relevant for the xerophytic lifestyle of conifers than a large water reservoir inside the endodermis even under fluctuating dry conditions or drought. Under fluctuating dry conditions, osmotic water uptake is still needed for phloem loading that already might commence in the pre-phloem pathway, utilising aquaporins in the transfusion parenchyma (Laur & Hacke, 2014; Schulz, 2015). In this situation, competition between the three water sinks in the needle - evaporation, photosynthesis and phloem loading - is controlled by the endodermis, and the apoplasmic domains of stele and mesophyll are uncoupled.

Under lasting drought conditions, the transfusion tissue contributes to cavitation repair (for review see (Brodersen & McElrone, 2013)). Cavitation can result from drought or freezing events in the winter season. One way to cope with it are the specific bordered pits between the axial xylem tracheids in needles and the wood (Hacke *et al*., 2015; Maruta *et al*., 2020). Another one was shown in *Taxus baccata*, where declines in hydraulic conductance were accompanied with the reversible deformation of the transfusion tracheid system (Zhang *et al*., 2014). When assessing ultrasonic acoustic emission as indicator for cavitation, Johnsen and co-workers found that - according to cryo-SEM - the transfusion tissue of needles emptied without acoustic signals, in contrast to the axial tracheid system (Johnson *et al*., 2009). This might be due to the fact that the thin transfusion tracheid walls do not resist compression (see Fig. 1 (Carde, 1978). The broad-leaved rainforest conifer *Podocarpus greyii* even has a specialised accessory transfusion tracheid system that showed strong deformation under water stress (Brodribb & Holbrook, 2005).

Sunken stomata and the small overall surface-to-volume ratio of conifer needles are one indication of a xerophytic lifestyle; the endodermis-type bundle sheath and the transfusion tissue together might be considered yet another specific adaption to cold and drought conditions. As in roots, the Casparian strip hermetically closes the stelar apoplast, as shown with the iohexol experiments: no iohexol left the stele, not even after 36-hour treatments, while the membrane permeable potassium iodide did so (Fig. 1). However, under desiccation stress this direction might be inverted for foliar water uptake to assist in refilling embolised xylem. The Casparian strip blocks the endodermis in either direction as shown by incubation of the wounded needle epidermis with the apoplast marker calcofluor (Wu *et al*., 2005). Water taken up via the needle surface and/or stomata will therefore cross the endodermis through its aquaporins (Mayr *et al*., 2014).

Cavitation repair is generally considered an osmotically driven process, involving phloem and rays in the stem (Brodersen & McElrone, 2013). The complex transfusion tissue with its intervening sugar-filled parenchyma and apoplasmic seems to be a needle adaptation to protect these evergreen organs in winter, dry conditions and drought. Depending on the physiological conditions, the transfusion tracheid system is an intrastelar water storage tissue which provides water not only outward to transpiration and photosynthesis, but also inward to phloem loading and cavitation repair. Like succulent plants shrink and swell depending on water availability, the transfusion tissue can shrink to cope with hydraulic fluctuations. These fluctuations can then osmotically be balanced by the living transfusion parenchyma which filled around 60% of the stele under well-watered conditions (Table 1). The transfusion tissue can also swell: freezing led in *Pinus radiata* needles to the increase in transfusion tracheid volume and ice crystal formation, while the transfusion parenchyma shrunk (Roden *et al*., 2009). Release of water from the parenchyma increased the freeze protection of the living tissue by increasing the sugar concentration. Synchrotron imaging of *P. pinaster* needles showed indeed that dehydration led to large changes in the volume of the transfusion tissue up to nearly 50 % at – 8 MPa and finally deformed the entire needle (Bouche *et al*., 2016).

The complexity of the transfusion tissue reflects its multiple functions under ever changing environmental conditions. It takes care of the outward transport of water from the xylem to the mesophyll for photosynthesis, and the inward transport of sugars from the mesophyll to the phloem. However, its structure also reveals an adaptation to efficiently regulate water allocation when the needle is exposed to fluctuating desiccation conditions and even drought. The key seems to be that both functions – dead water reservoir and living sugar pathway – are inseparable, but how the functions are adjusted transiently and permanently, sometimes even involving the inversion of the transport directions, remains to be addressed in future studies.

## Acknowledgements

We thank the Independent Research Fund Denmark for granting the project, the Center for Advanced Bioimaging, University of Copenhagen, for providing its electron and fluorescence microscopes, and the 3D Imaging Center at the Technical University of Denmark for access to the X-ray imaging instrument.

## Author contributions

AS, HFP and TB designed and supervised the study, ChG performed the anatomical and physiological experiments and prepared the plant material for μXCT, CaG developed the setup of the X-ray microscope and handled the imaging data, AS, ChG and SJVM segmented the imaging data, SJVM and TB analysed cell connectivity in the stele, ChG, SJVM AS and TB drafted the manuscript, and all authors contributed to discussion and interpretation of data and revision of the manuscript.

## Supporting Information

Additional supporting information may be found online in the Supporting Information section at the end of the article

**Fig. S1** Chamber used for μXCT scanning as 3D drawing and mounted in the imaging instrument

**Fig. S2** *P. pinaster* needles stained with the fluorescent dye Coriphosphine O at different positions along the needle

**Fig. S3** Tube-like cell chain from the same μXCT stack as Fig. 3, 3D-rendered as a 27 μm thick slice corresponding to Fig. 3c

**Fig. S4** Small air inclusions in the centre of the stele form fibre-like axial structures extending throughout the scanned volume

**Fig. S5** Same as Fig 1b but with a look-up-table accommodating colour-vision impaired readers **Video S1** Animation following all cross sections through a μXCT stack of a *P. pinaster* needle segmented for the iohexol-positive apoplast (magenta), iohexol-negative tissue (green), and air (blue).

**Video S2** Animation following the tangential sections through a μXCT stack of a *P. pinaster* needle segmented for the iohexol-positive apoplast (magenta), iohexol-negative tissue (green), and air (blue).

**Video S3** Animation of the 3D structure of the complex apoplast of a *P. pinaster* needle (same as Fig 3d)

## Supporting Information

**Fig. S1.**
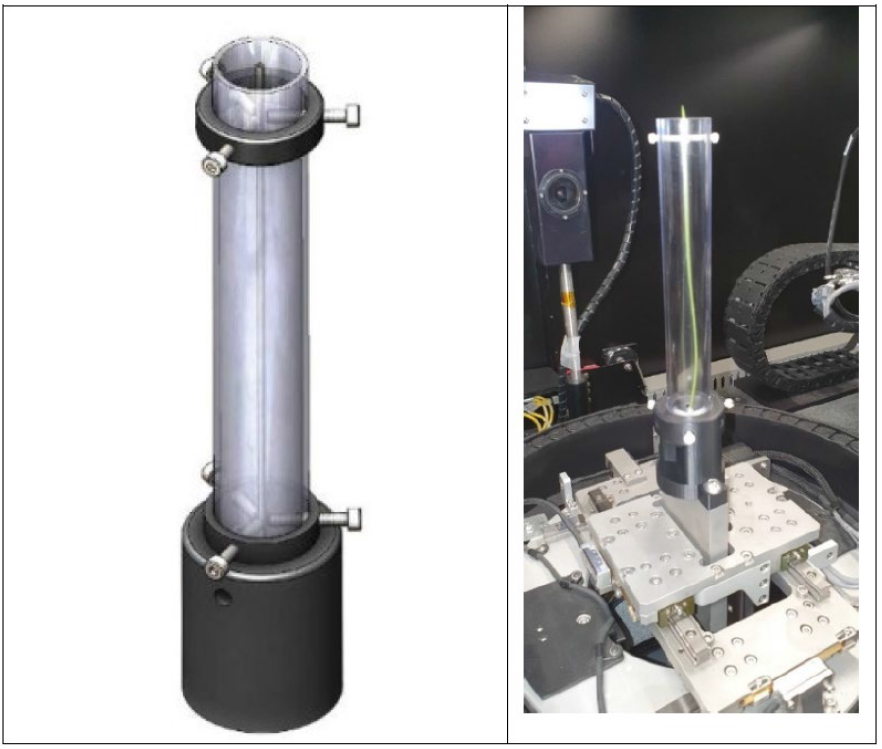
Chamber used for μXCT scanning as 3D drawing and mounted in the imaging instrument

**Fig. S2.**
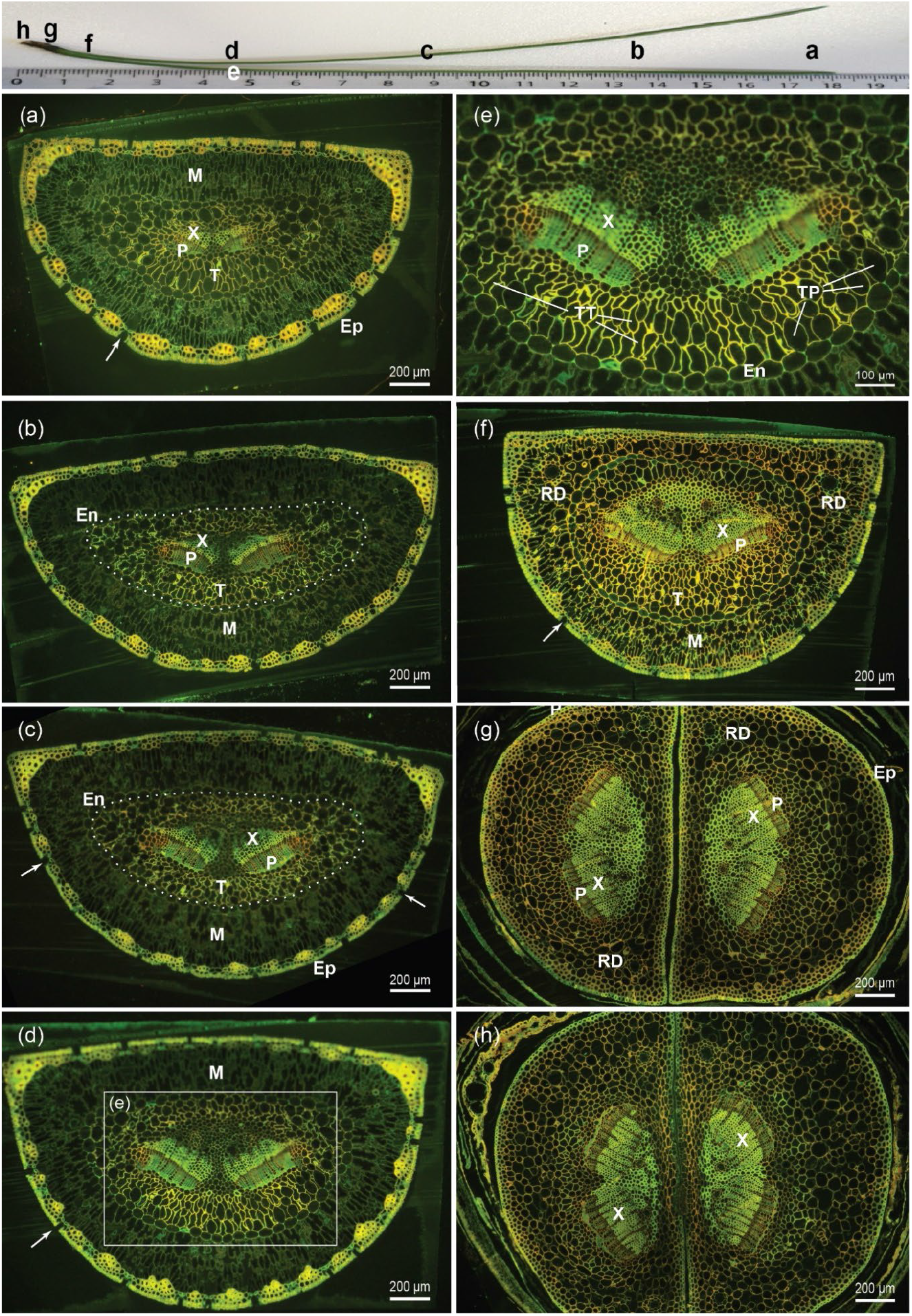
*P. pinaster* needles stained with the fluorescent dye Coriphosphine O at different positions along the needle as indicated in the top photograph. (a-f) Cross sections show two vascular bundles each with xylem (X) and phloem (P). Epidermis (Ep) with sunken stomata (arrows) borders hypodermal fibers and the mesophyll (M). The endodermis-type bundle sheath (En) is marked with a stippled line in (b) and (c). (e) Enlarged view of the stele at the boxed region of (d). The transfusion tissue inside the endodermis contains transfusion tracheids (TT) and transfusion parenchyma (TP), both abutting the endodermis. (g-h) Cross-sections of the fascicle base. The needle pairs are still closely connected to each other and surrounded by the fascicular sheath

**Fig. S3.**
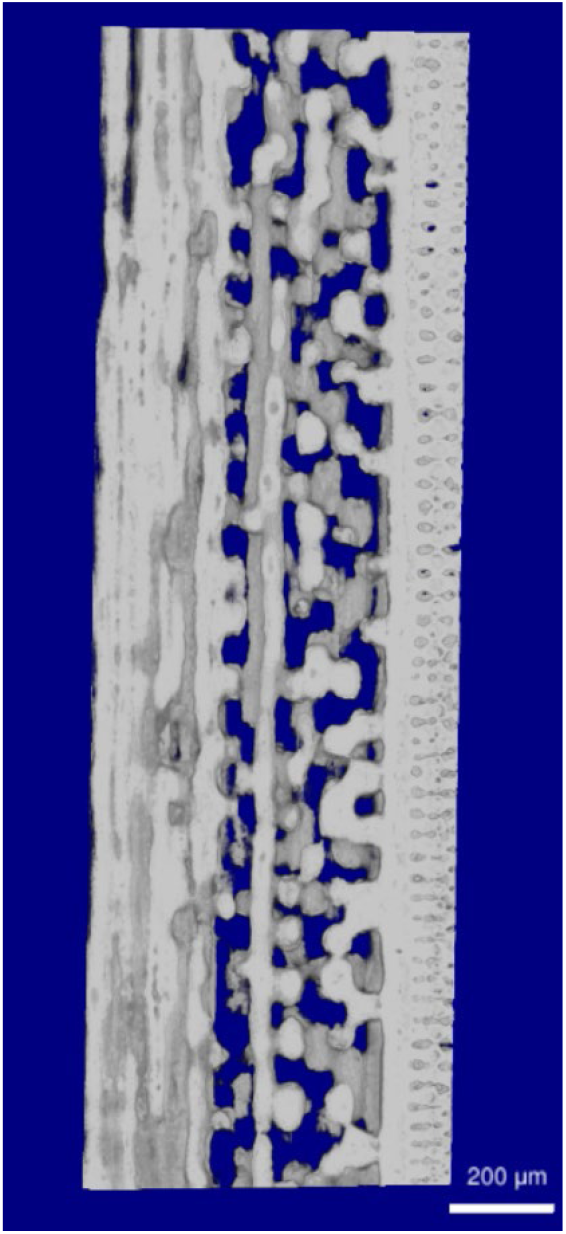
Tube-like cell chain from the same μXCT stack as Fig. 3, 3D-rendered as a 27 μm thick slice corresponding to Fig. 3c

**Fig. S4.**
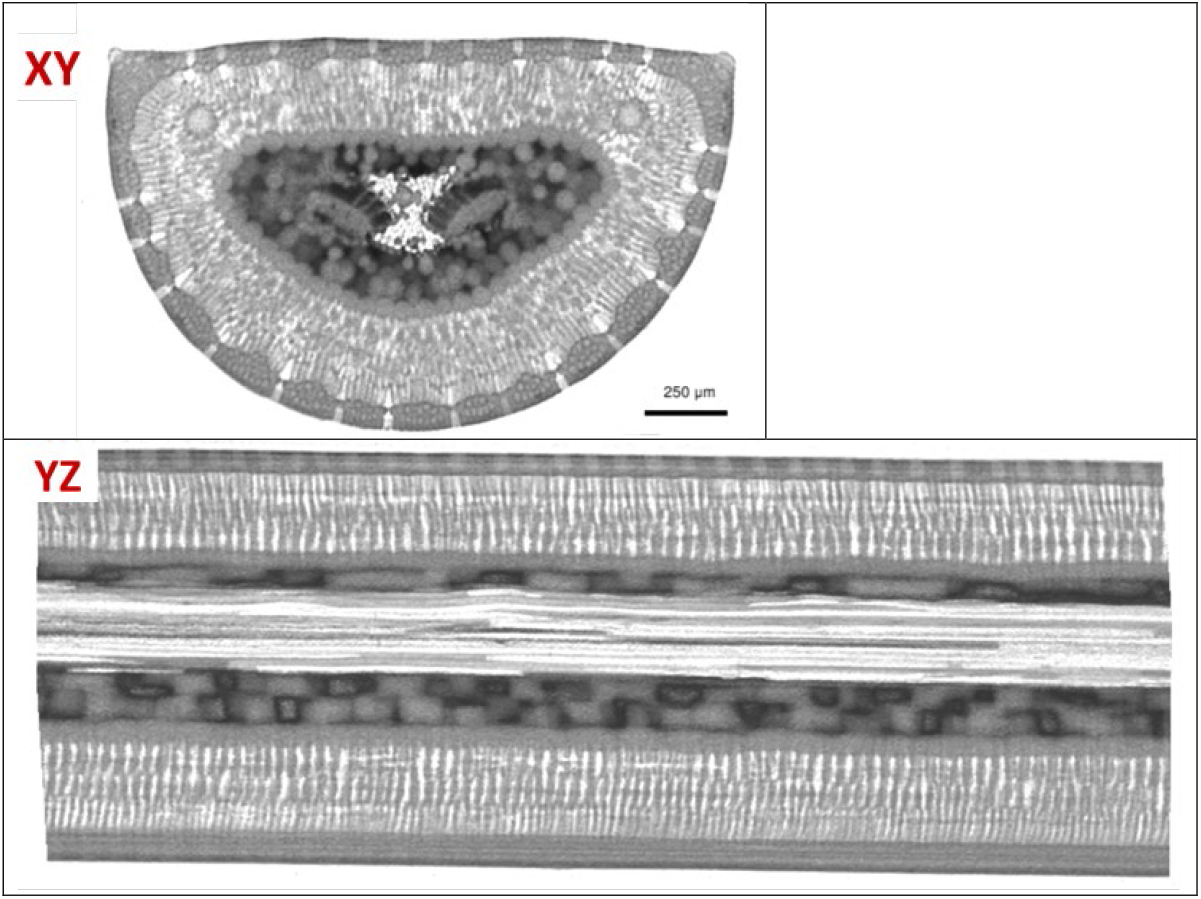
Needle cross section (XY); small air inclusions in the centre of the stele (white) contrast to the iohexol black apoplast and the grey tissue contrast. Needle longitudinal section (XZ); the air inclusions clearly form fibre-like axial structures extending throughout the scanned volume

**Fig. S5.**
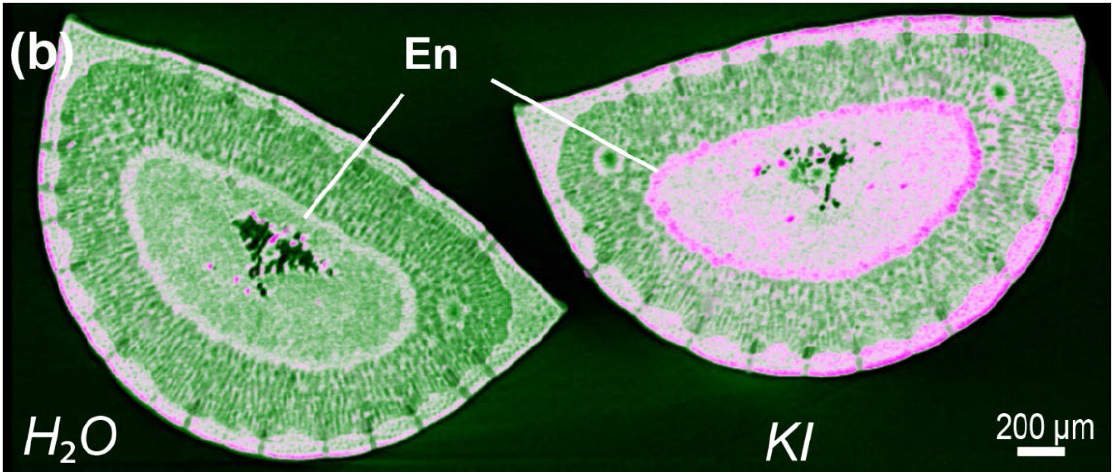
(Same as Fig 1b but with a look-up-table accommodating colour-vision impaired readers). Needle pair, supplied with water and potassium iodide, respectively and rendered as 50μm slice with a magenta-green look-up table to emphasise the KI accumulation in stele and endodermis (En) as well as mesophyll and epidermis

**Video S1** Animation following all cross sections through a μXCT stack of a *P. pinaster* needle segmented for the iohexol-positive apoplast (magenta), iohexol-negative tissue (green), and air (blue).

## References

Baird AS, Taylor SH, Pasquet-Kok J, Vuong C, Zhang Y, Watcharamongkol T, Scoffoni C, Edwards EJ, Christin PA, Osborne CP, et al. 2021. Developmental and biophysical determinants of grass leaf size worldwide. Nature 592(7853): 242–247.

Barberon M, Geldner N. 2014. Radial Transport of Nutrients: The Plant Root as a Polarized Epithelium. Plant Physiology 166(2): 528–537.

Barbour MM, Farquhar GD. 2004. Do pathways of water movement and leaf anatomical dimensions allow development of gradients in (H2O)-O-18 between veins and the sites of evaporation within leaves? Plant Cell and Environment 27(1): 107–121.

Bouche PS, Delzon S, Choat B, Badel E, Brodribb TJ, Burlett R, Cochard H, Charra-Vaskou K, Lavigne B, Li S, et al. 2016. Are needles of Pinus pinaster more vulnerable to xylem embolism than branches? New insights from X-ray computed tomography. Plant Cell and Environment 39(4): 860–870.

Brodersen CR, McElrone AJ. 2013. Maintenance of xylem network transport capacity: a review of embolism repair in vascular plants. Front Plant Sci 4.

Brodribb TJ, Feild TS, Jordan GJ. 2007. Leaf maximum photosynthetic rate and venation are linked by hydraulics. Plant Physiology 144(4): 1890–1898.

Brodribb TJ, Holbrook NM. 2005. Water stress deforms tracheids peripheral to the leaf vein of a tropical conifer. Plant Physiology 137(3): 1139–1146.

Brodribb TJ, Holbrook NM, Zwieniecki MA, Palma B. 2005. Leaf hydraulic capacity in ferns, conifers and angiosperms: impacts on photosynthetic maxima. New Phytologist 165(3): 839–846.

Buckley TN, John GP, Scoffoni C, Sack L. 2015. How Does Leaf Anatomy Influence Water Transport outside the Xylem? Plant Physiology 168(4): 1616–1635.

Canny MJ. 1993. Transfusion Tissue of Pine Needles as a Site of Retrieval of Solutes from the Transpiration Stream. New Phytologist 123(2): 227–232.

Carde J-P. 1978. Ultrastructural studies of *Pinus pinaster* needles: the endodermis. American Journal of Botany 65(10): 1041–1054.

Carde JP. 1973. Transfer Tissue (Strasburger Cells) in Needles of Maritime Pine (*Pinus pinaster* Ait).1. Histology and Ultrastructural Study of Grown up Tissue. Journal of Microscopy-Oxford 17(1): 65–88.

Carvalho MR, Losada JM, Niklas KJ. 2018. Phloem networks in leaves. Current Opinion in Plant Biology 43: 29–35.

Charra-Vaskou K, Mayr S. 2011. The hydraulic conductivity of the xylem in conifer needles (Picea abies and Pinus mugo). Journal of Experimental Botany 62(12): 4383–4390.

Couvreur V, Faget M, Lobet G, Javaux M, Chaumont F, Draye X. 2018. Going with the Flow: Multiscale Insights into the Composite Nature of Water Transport in Roots. Plant Physiology 178(4): 1689–1703.

Hacke UG, Lachenbruch B, Pittermann J, Mayr S, Domec J-C, Schulte PJ 2015. The Hydraulic Architecture of Conifers. In: Hacke U ed. Functional and Ecological Xylem Anatomy. Cham: Springer International Publishing, 39–75.

Han X, Turgeon R, Schulz A, Liesche J. 2019. Environmental conditions, not sugar export efficiency, limit the length of conifer leaves. Tree Physiol 39(2): 312–319.

Heimerdinger G. 1951. Zur Mikrotopographie der Saftströme im Transfusionsgewebe der Koniferennadel. Planta 40(2): 93–111.

Holmlund HI, Pratt RB, Jacobsen AL, Davis SD, Pittermann J. 2019. High-resolution computed tomography reveals dynamics of desiccation and rehydration in fern petioles of a desiccation-tolerant fern. New Phytologist 224(1): 97–105.

Huber B. 1947. ZUR MIKROTOPOGRAPHIE DER SAFTSTRÖME IM TRANSFUSIONSGEWEBE DERKONIFERENNADEL. I. Mitteilung. ANATOMISCHER TEIL. Planta 35: 331–351.

Humphrey OS, Young SD, Bailey EH, Crout NMJ, Ander EL, Hamilton EM, Watts MJ. 2019. Iodine uptake, storage and translocation mechanisms in spinach (Spinacia oleracea L.). Environ Geochem Health 41(5): 2145–2156.

Hunziker P, Schulz A 2019. Transmission electron microscopy of the phloem with minimal artefacts. In: Liesche J ed. Phloem: Methods and Protocols, Methods in Molecular Biology, vol. 2014: Springer, New York, 17–27.

Jensen KH, Berg-Sørensen K, Bruus H, Holbrook NM, Liesche J, Schulz A, Zwieniecki MA, Bohr T. 2016. Sap flow and sugar transport in plants. Reviews Modern Physics 88(035007): 1–63.

Jensen KH, Liesche J, Bohr T, Schulz A. 2012. Universality of phloem transport in seed plants. Plant Cell and Environment 35(6): 1065–1076.

Johnson DM, Meinzer FC, Woodruff DR, McCulloh KA. 2009. Leaf xylem embolism, detected acoustically and by cryo-SEM, corresponds to decreases in leaf hydraulic conductance in four evergreen species. Plant Cell and Environment 32(7): 828–836.

Kiferle C, Gonzali S, Holwerda H, Real Ibaceta R, Perata P. 2013. Tomato fruits: a good target for iodine biofortification. Front Plant Sci 4(205).

Laur J, Hacke UG. 2014. Exploring Picea glauca aquaporins in the context of needle water uptake and xylem refilling. New Phytologist 203(2): 388–400.

Liesche J, Martens HJ, Schulz A. 2011. Symplasmic transport and phloem loading in gymnosperm leaves. Protoplasma 248(1): 181–190.

Liesche J, Schulz A. 2012. In Vivo Quantification of Cell Coupling in Plants with Different Phloem-Loading Strategies. Plant Physiology 159(1): 355–365.

Liesche J, Vincent C, Han X, Zwieniecki M, Schulz A, Gao C, Bravard R, Marker S, Bohr T. 2021. The mechanism of sugar export from long conifer needles. New Phytologist 230(5): 1911–1924.

Maruta E, Kubota M, Ikeda T. 2020. Effects of xylem embolism on the winter survival of Abies veitchii shoots in an upper subalpine region of central Japan. Sci Rep 10(1): 6594.

Mayr S, Schmid P, Laur J, Rosner S, Charra-Vaskou K, Dämon B, Hacke UG. 2014. Uptake of Water via Branches Helps Timberline Conifers Refill Embolized Xylem in Late Winter Plant Physiology 164(4): 1731–1740.

McAdam SAM, Eleouet MP, Best M, Brodribb TJ, Murphy MC, Cook SD, Dalmais M, Dimitriou T, Gelinas-Marion A, Gill WM, et al. 2017. Linking Auxin with Photosynthetic Rate via Leaf Venation. Plant Physiology 175(1): 351–360.

Münch E. 1930. Die Stoffbewegungen in der Pflanze. Jena: Gustav Fischer.

Pongrac P, Baltrenaite E, Vavpetič P, Kelemen M, Kladnik A, Budič B, Vogel-Mikuš K, Regvar M, Baltrenas P, Pelicon P. 2019. Tissue-specific element profiles in Scots pine (Pinus sylvestris L.) needles. Trees 33(1): 91–101.

Pratt RB, Jacobsen AL. 2018. Identifying which conduits are moving water in woody plants: a new HRCT-based method. Tree Physiology 38(8): 1200–1212.

Rademaker H, Zwieniecki MA, Bohr T, Jensen KH. 2017. Sugar export limits size of conifer needles. Phys Rev E Stat Nonlin Soft Matter Phys 95(4-1): 042402.

Roden JS, Canny MJ, Huang CX, Ball MC. 2009. Frost tolerance and ice formation in Pinus radiata needles: ice management by the endodermis and transfusion tissues. Functional Plant Biology 36(2): 180–189.

Ronellenfitsch H, Liesche J, Jensen KH, Holbrook NM, Schulz A, Katifori E. 2015. Scaling of phloem structure and optimality of photoassimilate transport in conifer needles. Proc Biol Sci 282(1801): 20141863.

Sakurai G, Miklavcic SJ. 2021. On the Efficacy of Water Transport in Leaves. A Coupled Xylem-Phloem Model of Water and Solute Transport. Front Plant Sci 12: 615457.

Schulte PJ, Hacke UG. 2021. Solid mechanics of the torus–margo in conifer intertracheid bordered pits. New Phytologist 229(3): 1431–1439.

Schulz A 1990. Conifers. In: Behnke HD, Sjolund RD eds. Sieve Elements: Springer Berlin Heidelberg, 63–88.

Schulz A. 2015. Diffusion or bulk flow: how plasmodesmata facilitate pre-phloem transport of assimilates. Journal of Plant Research 128(1): 49–61.

Strasburger E 1891. Über den Bau und die Verrichtungen der Leitungsbahnen in den Pflanzen. Histologische Beiträge. Jena: Gustav Fischer.

Wu X, Lin J, Lin Q, Wang J, Schreiber L. 2005. Casparian Strips in Needles are More Solute Permeable than Endodermal Transport Barriers in Roots of Pinus bungeana. Plant and Cell Physiology 46(11): 1799–1808.

Zhang Y-J, Rockwell FE, Wheeler JK, Holbrook NM. 2014. Reversible Deformation of Transfusion Tracheids in Taxus baccata Is Associated with a Reversible Decrease in Leaf Hydraulic Conductance. Plant Physiology 165(4): 1557–1565.

Zwieniecki MA, Stone HA, Leigh A, Boyce CK, Holbrook NM. 2006. Hydraulic design of pine needles: one-dimensional optimization for single-vein leaves. Plant Cell and Environment 29(5): 803–809.

